# Naturalistic spatiotemporal stimulus modulation during epiretinal stimulation increases the persistence of retinal ganglion cell responsivity

**DOI:** 10.1101/2020.11.18.386961

**Authors:** Naïg Aurélia Ludmilla Chenais, Marta Jole Ildelfonsa Airaghi Leccardi, Diego Ghezzi

## Abstract

**Objective:** Retinal stimulation in blind patients evokes the sensation of discrete points of light called phosphenes, which allows them performing visual guided tasks, such as orientation, navigation, object recognition, object manipulation and reading. However, the clinical benefit of artificial vision in profoundly blind patients is still tenuous, as several engineering and biophysical obstacles keep it away from natural perception. The relative preservation of the inner retinal neurons in hereditary degenerative retinal diseases, such as retinitis pigmentosa, supports artificial vision through the network-mediated stimulation of retinal ganglion cells. However, the response of retinal ganglion cells to repeated electrical stimulation rapidly declines, primarily because of the intrinsic desensitisation of their excitatory network. In patients, upon repetitive stimulation, phosphenes fade out in less than half of a second, which drastically limits the understanding of the percept.

**Approach:** A more naturalistic stimulation strategy, based on spatiotemporal modulation of electric pulses, could overcome the desensitisation of retinal ganglion cells. To investigate this hypothesis, we performed network-mediated epiretinal stimulations paired to electrophysiological recordings in retinas explanted from both male and female retinal degeneration 10 mice.

**Main results:** The results showed that the spatial and temporal modulation of the network-mediated epiretinal stimulation prolonged the responsivity of retinal ganglion cells from 400 ms up to 4.2 s.

**Significance:** A time-varied, non-stationary and interrupted stimulation of the retinal network, mimicking involuntary microsaccades, might reduce the fading of the visual percept and improve the clinical efficacy of retinal implants.

## Introduction

The first notion of electrically-evoked visual percept came in 1755 when Charles Le Roy’s patient had reported seeing flashes of light while receiving an electric current across the head [1]. Almost three centuries later, three retinal stimulation systems (Argus^®^ II, Second Sight Medical Products; Alpha IMS and Alpha AMS, Retinal Implant AG) received marketing approval in Europe or United States of America for retinitis pigmentosa, a set of inherited retinal dystrophies, causing the progressive loss of retinal photoreceptors, the constriction of the visual field and eventually blindness [2].

The relative preservation of the inner retinal neurons in retinitis pigmentosa [3–5] supports artificial vision through the network-mediated stimulation of retinal ganglion cells (RGCs) either from the subretinal or the epiretinal side [6,7]. In epiretinal stimulation, RGCs can be stimulated either through direct activation of their axons and somas [8–10] or through indirect stimulation of their presynaptic neurons [7,11]. The latter strategy offers two advantages: it avoids the uncontrolled activation of distal axons leading to streak-like elongated phosphenes [12,13] and it provides spiking patterns resembling the natural response to visual stimuli [6,14]. However, the rapid desensitisation of the RGC response under repeated indirect stimulation is a significant limitation of this approach.

In animal models, the responsivity of RGCs to electrical stimulation rapidly decreases with repeated pulsed stimulation [15,16]. This decrease is likely caused by the intrinsic desensitisation of bipolar cells (BCs) [17], although an active contribution from the inhibitory network also has to be considered [16,17]. Moreover, the reduction of responsivity in RGCs might be associated with the phosphene fading reported by implanted patients during the clinical trials with the Argus^®^ II and the Alpha IMS retinal prostheses [18–20]. The time course of the RGC response reduction starts with a rapid loss of responsivity immediately after the first electric stimulus, and a slower response decay lasting up to several seconds follows for prolonged stimulation. This time course remarkably matches the two-phase decay in phosphene brightness reported by implanted patients under stationary vision [16].

In natural vision, during prolonged fixation, microsaccades counterbalance the transience of the retinal response [21]. Similarly, in retinal prostheses, a temporal and spatial modulation of the electric pulses may counteract the desensitisation process occurring because of the RGC network-mediated stimulation. Electric stimuli with randomised inter-pulse intervals induced in-vivo stronger electrically evoked potentials than regularly spaced pulses, thus suggesting the ability of temporal modulation to reduce the fading process [22]. Besides, in a study with high-density epiretinal indirect stimulation, we showed that alternating the stimulation among two neighbouring pixels allowed for the recovery of the RGC response at the pixel switch [23]. This result suggests that a RGC could be activated successively but independently through a different portions of its inner retinal network. This last property offers excellent potential to mimic the effect of microsaccades artificially, and it indicates the use of spatial modulation to reduce the fading process. Therefore, we investigate whether a more naturalistic stimulation with spatiotemporal modulations, including time-varied, non-stationary and interrupted features can increase the persistence of the prosthetic responses at the RGC level.

## Materials and Methods

### POLYRETINA micro-fabrication

Photovoltaic interfaces were fabricated on silicon wafers. A thin sacrificial layer of poly(4-styrenesulfonic acid) solution (561223, Sigma-Aldrich) was spin-coated on the wafers (1500 rpm, 40 s) and baked (135 °C, 10 min). Degassed PDMS pre-polymer (10:1 ratio base-to-curing agent, Sylgard 184, Dow-Corning) was then spin-coated (500 rpm, 60 s) and cured in the oven (80 °C, 2 hr). PEDOT:PSS (PH1000, Clevios Heraeus) was mixed to 0.1 v/v% (3-glycidyloxypropyl)trimethoxysilane (440167, Sigma Aldrich), ultra-sonicated for 20 min, filtered (0.2 μm PES filters), and then spin-coated at 3000 rpm for 40 seconds. Subsequent annealing at 115 °C for 30 min was performed. The preparation of the bulk heterojunction was performed in a glove box under nitrogen atmosphere. 20 mg of P3HT (M1011, Ossila) and 20 mg of PC_60_BM (M111, Ossila) were dissolved in 1 mL of anhydrous chlorobenzene each and let stirring overnight (16 hr) at 70 °C. The solutions were then filtered (0.45 μm PTFE filters) and blended (1:1 v:v). The P3HT:PC_60_BM blend was spin-coated at 1000 rpm for 45 seconds. Subsequent annealing at 115 °C for 30 min was performed. Titanium and titanium nitride were deposited respectively by direct-current and radio frequency magnetron sputtering using a shadow mask. Polymers were patterned by oxygen plasma. The wafers were then placed in deionised water to allow for the dissolution of the sacrificial layer and the release of the photovoltaic interfaces. The floating membranes were finally collected and dried in air.

### Preparation of retinal explants

Animal experiments were conducted according to the animal authorisation GE3717 issued by the Département de l’Emploi, des Affaires sociales et de la Santé (DEAS), Direction Générale de la Santé of the République et Canton de Genève (Switzerland). Retinal degeneration 10 (rd10) mice (aged 121.5 ± 13.8 days; mean ± s.d.) were sacrificed under normal light conditions by injection of sodium pentobarbital (150 mg/kg), and eyes were immediately collected and dissected in carboxygenated (95% O2 and 5% CO_2_) Ames’ medium (A1420, Sigma-Aldrich). Whole-mount retinas were placed ganglion cell down onto the stimulation pixels and transferred to the microscope stage for stimulation and recording. During the entire preparation and recording procedures, the retinas were maintained under dim red light and perfused with carboxygenated Ames’ medium at 32 °C.

### Electrophysiological recordings

Extracellular recordings were performed with a sharp metal electrode (PTM23BO5KT, WPI), amplified (Model 3000, A-M System), filtered (300-3,000 Hz) and digitalized at 30 kHz (Micro1401-3, CED). RGCs were identified by their spontaneous spiking activity. Spike detection and spike sorting were performed offline by threshold detection using the wave_clus algorithm [24]. Data processing was performed in MATLAB (MathWorks).

### Retina stimulation

Retinal explants were stimulated with the POLYRETINA photovoltaic interface. 80-μm photovoltaic pixels separated by a 120-μm pitch were individually illuminated from a Nikon Ti-E inverted microscope (Nikon Instruments) using a Spectra X illumination system (Emission filter 560/32, Lumencor). The microscope was equipped with a dichroic filter (FF875-Di01-25×36, Semrock) and a 10x (CFI Plan Apochromat Lambda) objective. The timing, spatial patterning and alternation of the light stimuli were carried out with a light patterning system (Polygon 400, Mightex). Each light pattern was aligned to the pixels in real-time. Green light (560-nm) was projected with a 0.9 mW/mm^2^ irradiance for all the stimuli.

### Experimental Design and statistical analyses

The experimental design is shown in Tab. 1. Statistical analysis and graphical representation were performed in MATLAB. For each stimulation pulse, the network-mediated medium-latency (ML) response was calculated as the average firing rate elicited in a ± 20-ms window around the highest bin of the peri-stimulus time histogram (PSTH). The highest bin was screened for from 40 to 120 ms after the stimulus onset. The D’Agostino & Pearson omnibus normality test was performed to justify the use of a parametric or non-parametric test. The Tukey’s honestly significant difference post-hoc test was used to verify the use of multiple comparison tests.

**Table 1:** Animals and experimental conditions.

| Experimental group | Number of animals | Number of RGCs | Post-natal days (mean ± s.d.) | Females / Males | Stimulation protocols |
| --- | --- | --- | --- | --- | --- |
| A | 4 | 16 | 121.7 ± 10.6 | 2/2 | Figure 1 and Figure 2 |
| B | 3 | 22 | 113.0 ± 8.7 | 0/3 | Figure 3 |
| C | 3 | 12 | 115.3 ± 12.7 | 0/4 | Figure 4 and Figure 5 |
| D | 4 | 23 | 132.2 ± 17.1 | 2/2 | Figure 6 and Figure 7 |

### Data availability

The authors declare that all the relevant data supporting their findings are available in this article. Access to the raw data can be obtained from the corresponding author upon reasonable request.

## Results

### Frequency dependent desensitisations of RGCs

Epiretinal indirect activation of RGCs is subjected to strong desensitisation under repeated electrical stimulation because of its network-mediated mechanism. First, we evaluated the response decay of RGCs from explanted rd10 retinas (group A in Tab. 1) elicited by 10-ms photovoltaic stimulation from a selected pixel (stationary) repeated at 1 Hz, 5 Hz, 10 Hz and 20 Hz for 10 s using the POLYRETINA epiretinal prosthesis (Fig. 1A). In agreement with previous studies [15,25,16,26], we found that RGCs showed a strong naïve spiking response to the first pulse in the train (Fig. 1B,C) reaching firings rate up to 230 Hz. This response was significantly higher than the corresponding resting activity for each of the stimulation rates tested (p < 0.0001, p = 0.0001, p = 0.0069 and p = 0.0016 respectively for 1, 5, 10 and 20 Hz stimulation rate, two-tailed paired t-test). After 10 s of stimulation (Fig. 1C), only the 1-Hz stimulation rate evoked network-mediated ML responses significantly higher than the corresponding resting activity (p = 0.0030, two-tailed paired t-test). For higher stimulation rates, the response was not significantly higher than the corresponding resting activity anymore after 1.8, 0.4 and 0.25 s respectively for 5, 10 and 20 Hz stimulation rates (p = 0.4488, p = 0.0642 and p = 0.1831 respectively, two-tailed paired t-test).

**Figure 1.**
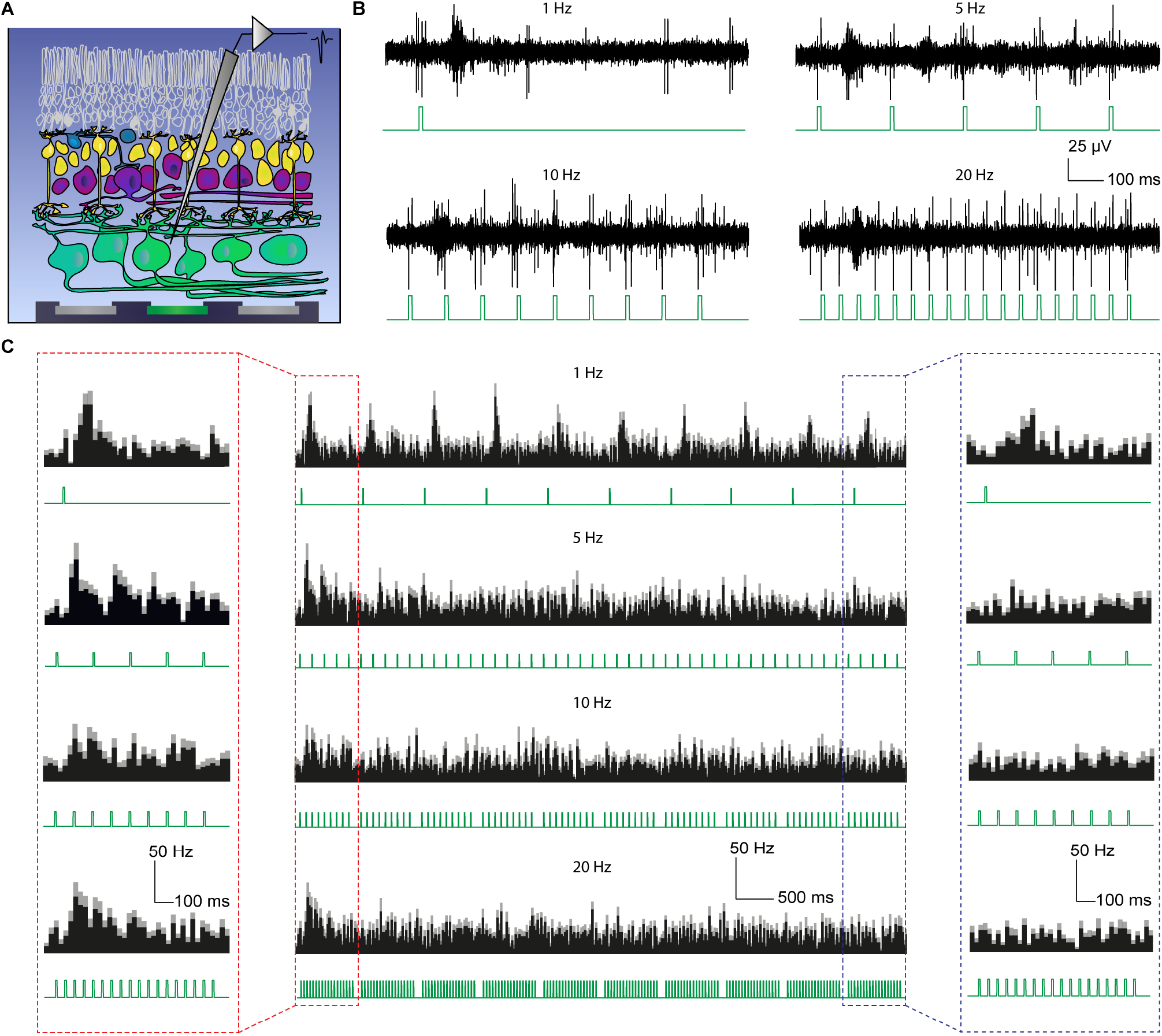
RGC responses to repeated indirect photovoltaic stimulation. **A**, Sketch of the stimulation and recording set-up. A single pixel (in green) of the POLYRETINA device was illuminated with 10-ms pulses (560 nm, 0.9 mW/mm^2^). Solid and empty cells represent respectively preserved (RGCs, ACs, BPs and horizontal cells) and degenerated (photoreceptors) cells in the rd10 mice model. **B**, Representative recordings of a RGC response to single-pixel illumination at 1, 5, 10 and 20 Hz for 1 s. The photovoltaic light pulses are shown in green. **C**, Mean PSTHs upon 10 consecutive sweeps of 1 s (10 s in total) at 1, 5, 10 and 20 Hz (mean ± s.e.m., group A, n = 16 RGCs). The filled area indicates the mean value, while the grey area shows the s.e.m. The first and tenth sweeps are highlighted respectively in red and blue. The corresponding light pulses are shown in green.

The network-mediated ML response rapidly decayed already from the second pulse of the sequence (Fig. 2A). We computed the response persistence, as the time between the first pulse in the train sequence and the first pulse evoking a network-mediated ML response statistically significantly lower (at least p < 0.05) than the naïve response to the first pulse. With 1-Hz stimulation rate (Fig. 2A, green), the network-mediated ML response became statistically significantly lower after 3 s (fourth pulse in the sequence; p = 0.038, two-tailed paired t-test). The response decay became faster as the stimulation rate increased: with 5-Hz stimulation rate (Fig. 2A, blue) the response persisted for 0.4 s (third pulse in the sequence; p = 0.0278, two-tailed paired t-test), at 10-Hz stimulation rate (Fig. 2A, red) for 0.5 s (sixth pulse in the sequence; p = 0.0183, two-tailed paired t-test) and at 20-Hz (Fig. 2A, orange) for 0.25 s (sixth pulse in the sequence; p = 0.0329, two-tailed paired t-test).

**Figure 2.**
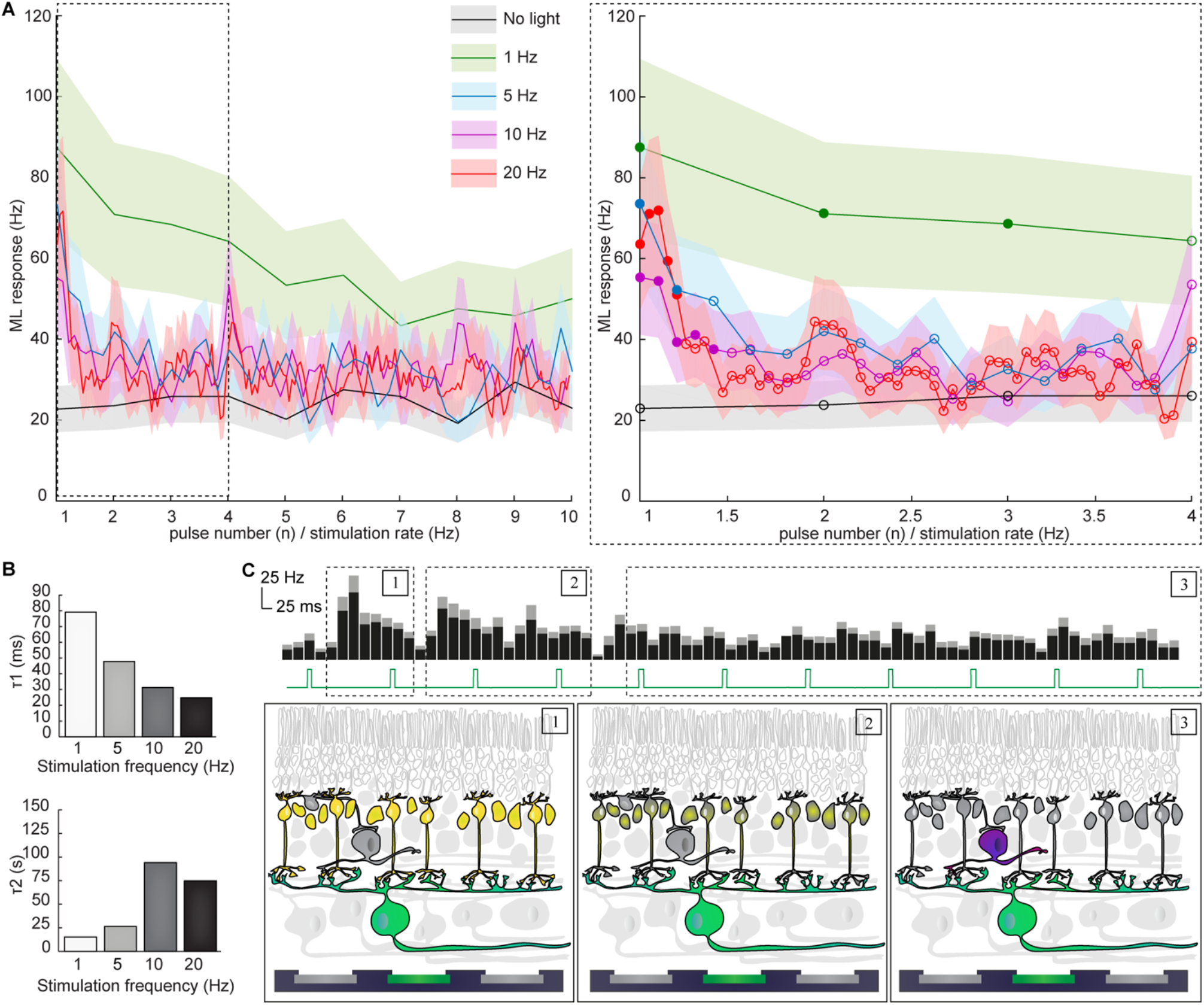
Double exponential decay of network-mediated ML RGC activity. **A**, Quantification of the network-mediated ML RGC response over 10 s of repetitive stimulation at 1, 5, 10 and 20 Hz and without light stimulation (mean ± s.e.m., group A, n = 16 RGCs). The right panel highlights the early desensitisation up to 2 s from the start of the pulse train. The shaded area shows the s.e.m. Filled circles represent pulses inducing a network-mediated ML activity not statistically significantly different than the naïve repose to the first pulse. Empty circles represent pulses inducing a network-mediated ML activity statistically significantly reduced compared to the naïve repose to the first pulse. **B**, Time constants of the fast (τ1) and slow (τ2) desensitisation components, extracted from the double exponential model for 1, 5, 10 and 20 Hz stimulation rates. **C**, Mechanism of excitation-inhibition balance chronology during RGC desensitisation. Solid and empty cells represent respectively preserved and degenerated cells in the rd10 mice model. The highlighted cells show the recorded RGC and its presynaptic neurons, among which resting cells are shown in grey, and depolarised cells are coloured. Panels illustrate the naïve network response (1), the early desensitisation phase (2) and the later desensitised state (3) respectively. The illustrative PSTH corresponds to a 5-Hz stimulation rate (2.5 s; mean ± s.e.m., group A, n = 16 RGCs). The filled area indicates the mean value, while the grey area shows the s.e.m.

The quantification of the network-mediated ML response decay revealed that it was inversely proportional to the stimulation frequency (Fig. 2A,B). In agreement with previous hypotheses describing a two-phases process behind RGCs desensitisation [16], the decay curves were fitted with a double exponential model. For all the stimulation rates tested, we found a first rapid phase occurring immediately after the first stimulation pulse, and a slower phase dominating the decay during prolonged stimulations. The decay rate of the rapid phase ranged from 25.6 ms to 79.0 ms (for 1 Hz and 20 Hz respectively), and it exponentially decreased with the increase of the stimulation frequency (R^2^ = 0.99; SSE = 3.45.10^-6^). The decay rate of the slow phase ranged from 19.7 ms to 93.7 ms, but it did not correlate to the stimulation frequency (Fig. 2B).

It was previously proposed that the rapid decay reflected the intrinsic desensitisation of the excitatory network [16,26] since the BCs in the rd10 retina were strongly desensitised upon electrical stimulation with frequencies above 4-6 Hz. Also, for stimulation rates exceeding 14 Hz, a repolarisation period with a time constant of 39.49 ms per pulse was reported to be necessary for the BC repolarisation [17]. Nevertheless, the involvement of the inhibitory network in the RGC desensitisation process could not be formally excluded [16,17]. Amacrine cells (ACs) are likely to play an active role in the RGC response during epiretinal stimulation [13], as well as to be the source of the slow desensitisation phase [16,26]. In this perspective, the two-phases RGC desensitisation would reflect the excitatory-inhibitory activity balance of the retinal network (Fig 2C): at the first pulse, the naïve BCs strongly depolarise and generate a high-frequency burst in the targeted RGCs (1); since BCs rapidly lose their intrinsic excitability, their input to RGCs in response to the next pulses is continuously decreased (2), and the inhibition from ACs becomes predominant (3).

### Temporal modulation of epiretinal stimulation with irregular pulse trains

The inverse relation between the fast decay rate and the stimulation frequency underlies the relevance of a recovery period between pulses. Increasing the inter-pulse interval could allow the recovery of excitability in BCs and reduce the desensitisation process. However, an increase in the inter-pulse interval does not necessarily imply to lower the stimulation frequency, since it can be achieved using a time-varying stimulation train in which each stimulation pulse has a different pulse duration. During natural vision, the light sequence reaching a steady location on the retina exhibits a highly irregular temporal profile, due to both eye and object movements. This continuous dynamic change contributes to counteract the desensitisation of BCs to natural stimuli [27,28]. Likewise, the use of time-varied and amplitude-varied pulse trains was shown to reduce fading of electrically evoked potentials recorded in the superior colliculus of healthy rats [22].

We tested randomised time-varying pulses (temporal modulation) in explanted rd10 retinas (group B in Tab. 1) upon photovoltaic single-pixel illumination at 5-Hz repetition rate (Fig. 3). The stimulation rate was set to 5 Hz as it was shown to be a comfortable stimulation rate for most patients [26,29], while it is low enough to allow the temporal modulation of the electric pulses. Pulse durations were varied between 5 ms and 25 ms, and random sequences were delivered for 10 s. Each RGC was tested with a different random sequence, whose cumulative exposure time was kept equal to 50 ms every 5 pulses for a total of 500 ms for the whole 10-s protocol, as for the 10-ms stationary stimulation. The fast response decay was slowed down by randomised time-varying stationary pulses (Fig. 3A,B), which allowed a network-mediated ML response persistence up to 0.8 s (Fig. 3C): the network-mediated ML response becoming statistically significantly lower than the naïve response from the fifth pulse in the sequence (p = 0.0154, two-tailed paired t-test). On the other hand, the response to 10-ms stationary pulses persisted only for 0.4 s (third pulse in the sequence; p = 0.0255, two-tailed paired t-test), as in the previous experiment.

**Figure 3.**
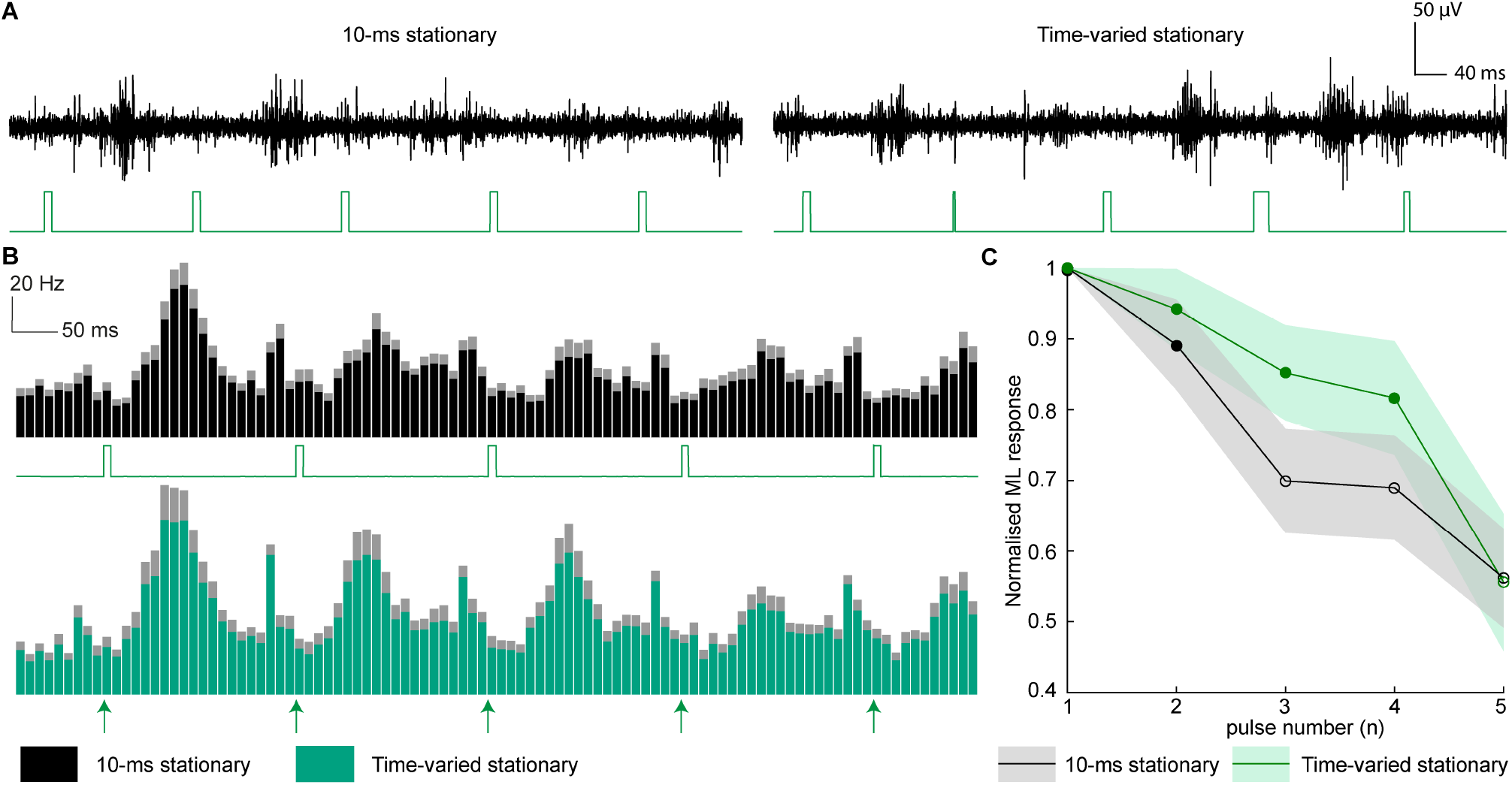
Randomised time-varied stationary pulse trains reduced RGC desensitisation. **A**, Representative recordings from a RGC upon 5-Hz single-pixel photovoltaic stimulation (5 light pulses for 1 s) with 10-ms stationary pulses (left) and randomised time-varying stationary pulses (right). The light pulses are shown in green. **B**, Mean PSTHs (mean ± s.e.m., group B, n = 22 RGCs) upon 5-Hz single-pixel photovoltaic stimulation (5 light pulses for 1 s) with 10-ms stationary pulses (top) and randomised time-varying stationary pulses (bottom). The filled area indicates the mean value, while the contoured area shows the s.e.m. Because of the randomly varying duration of each pulse over sequences, only the onset of the light pulses is indicated by the green arrows. **C**, Quantification of the normalised network-mediated ML RGC response over 1 s of repetitive stimulation at 5 Hz (mean ± s.e.m., group B, n = 22 RGCs) with 10-ms stationary pulses (black) and randomised time-varying stationary pulses (green). The firing rates were normalised to the naïve response to the first pulse. The shaded area shows the s.e.m. Filled circles represent pulses inducing a network-mediated ML activity not statistically significantly different than the naïve repose to the first pulse. Empty circles represent pulses inducing a network-mediated ML activity statistically significantly reduced compared to the naïve repose to the first pulse.

In summary, pulse trains with irregular pulse durations showed a transient ability to reduce the RGC desensitisation upon repeated electrical stimulation, lengthening the RGC response from 0.4 s to 0.8 s.

### Spatial modulation of epiretinal stimulation with non-stationary pulse trains

Another strategy exploited in natural vision to reduce fading is performing a great variety of eye movements during visual tasks. Saccades, eye drifts and microsaccades allow focusing on a specific object, while slightly shifting it to create a non-stationary image projection on the retina [30–32]. A previous experiment with singlepixel photovoltaic stimulation in rd10 retinas using the high-density POLYRETINA device suggested that non-stationary stimulation of the retina can be achieved by alternating the stimulation via neighbouring pixels, provided that the electrode density is high enough for more than one pixel to activate the same RGC [23]. The POLYRETINA prosthesis can indeed generate highly focused stimuli from each of its independent pixels, and because of its high resolution, a single RGC can be stimulated through two adjacent pixels.

Thus, we tested whether mimicking microsaccades with an alternating stimulation pattern (spatial modulation) in which the stimulation is switched between two adjacent pixels (10-ms non-stationary pulses) could reduce the RGC response desensitisation (Fig. 4). After the identification of a RGC and its receptive field (group C in Tab. 1), the illumination spot was switched at a 1-Hz rate between the two most responding pixels of its receptive field (10-ms pulses, 5-Hz stimulation rate). With 10-ms stationary stimulation, the network-mediated ML activity is reduced due to the desensitisation process (Fig. 4A left and 4B top), and the response persisted only up to 0.4 s, as before (Fig. 5A, black). When the stimulation is switched to the neighbouring pixel (Fig. 4A middle and 4B middle), the network-mediated ML activity is transiently recovered (Fig. 5A green), and it is not statistically significantly lower than the naïve response to the first pulse for two consecutive pulses (pulse 6: p = 0.9003, two-tailed paired t-test; pulse 7: p = 0.0835, two-tailed paired t-test). The stimulation from each of the two alternating pixel evoked persisting responses, lengthening the total RGC response duration up to 1.4 s (Fig. 5A). The pixel switch experiment was repeated with randomised timevarying non-stationary pulses, and identical results were obtained (Fig. 4A right, 4B bottom and 5A orange). However, when the stimulation was switched back to the first pixel (second switch, pulse number 11), the network-mediated ML activity was statistically significantly reduced compared to the naïve response for both 10-ms non-stationary (p = 0.0271, two-tailed paired t-test) and randomised time-varying non-stationary pulses (p = 0.0267, two-tailed paired t-test).

**Figure 4.**
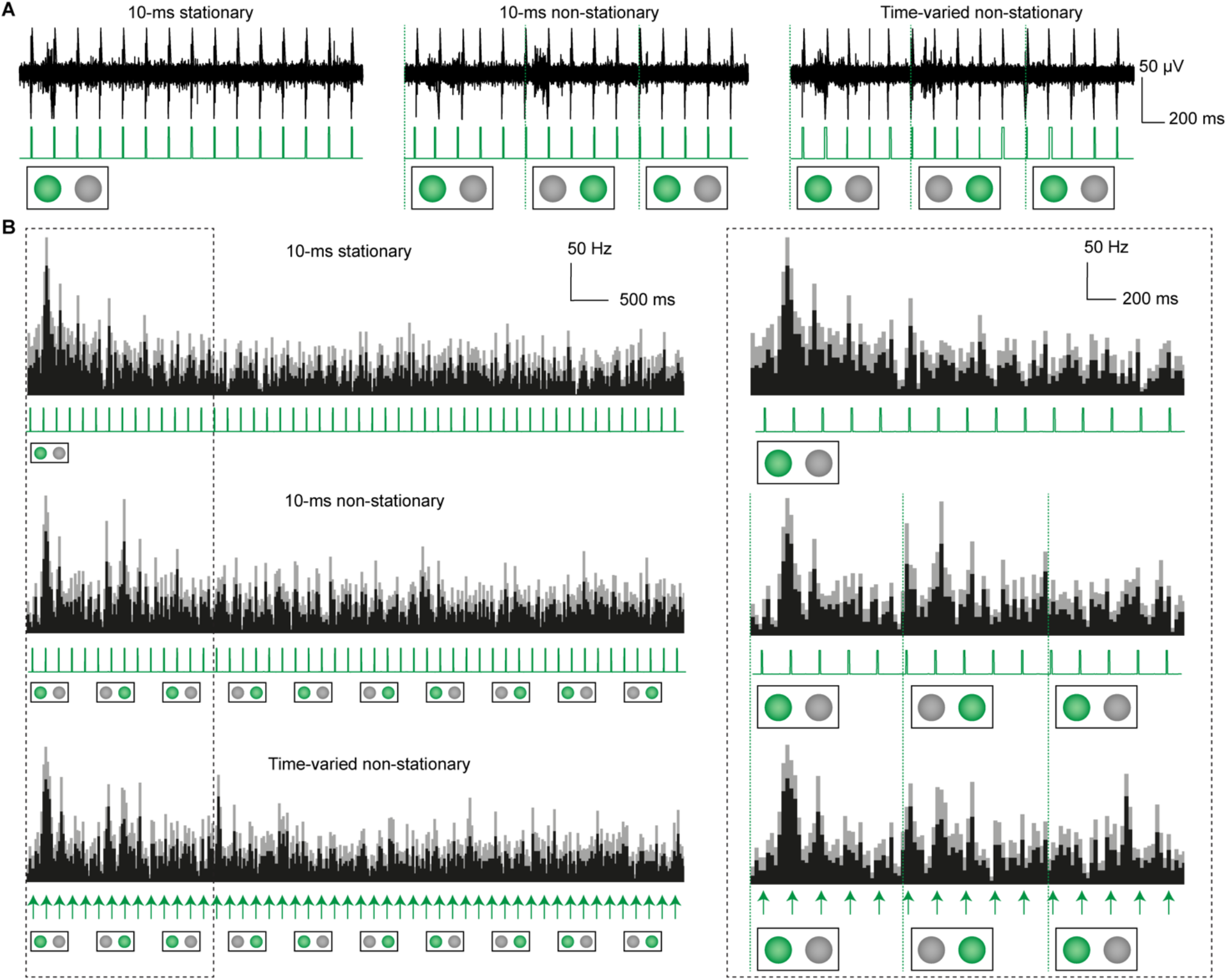
Non-stationary pulse trains reduced RGC desensitisation. **A**, Representative recordings from a RGC upon 5-Hz single-pixel photovoltaic stimulation (15 light pulses for 3 s) with 10-ms stationary pulses (left), 10-ms non-stationary pulses (middle) and randomised time-varying non-stationary pulses (right). The light pulses are shown in green. For non-stationary stimulation, the stimulating pixels were switched at 1 Hz. **B**, Mean PSTHs (mean ± s.e.m., group C, n = 12 RGCs) upon 5-Hz single-pixel photovoltaic stimulation (15 light pulses for 3 s) with 10-ms stationary pulses (top), 10-ms non-stationary pulses (middle) and randomised time-varying non-stationary pulses (bottom). The black area shows the mean value, while the grey area shows the s.e.m. The right panels highlight the first 3 s of the recordings. The light stimuli are indicated in green (top and middle), while the onset of the light pulses is indicated with green arrows for the randomised time-varying pulses (bottom).

**Figure 5.**
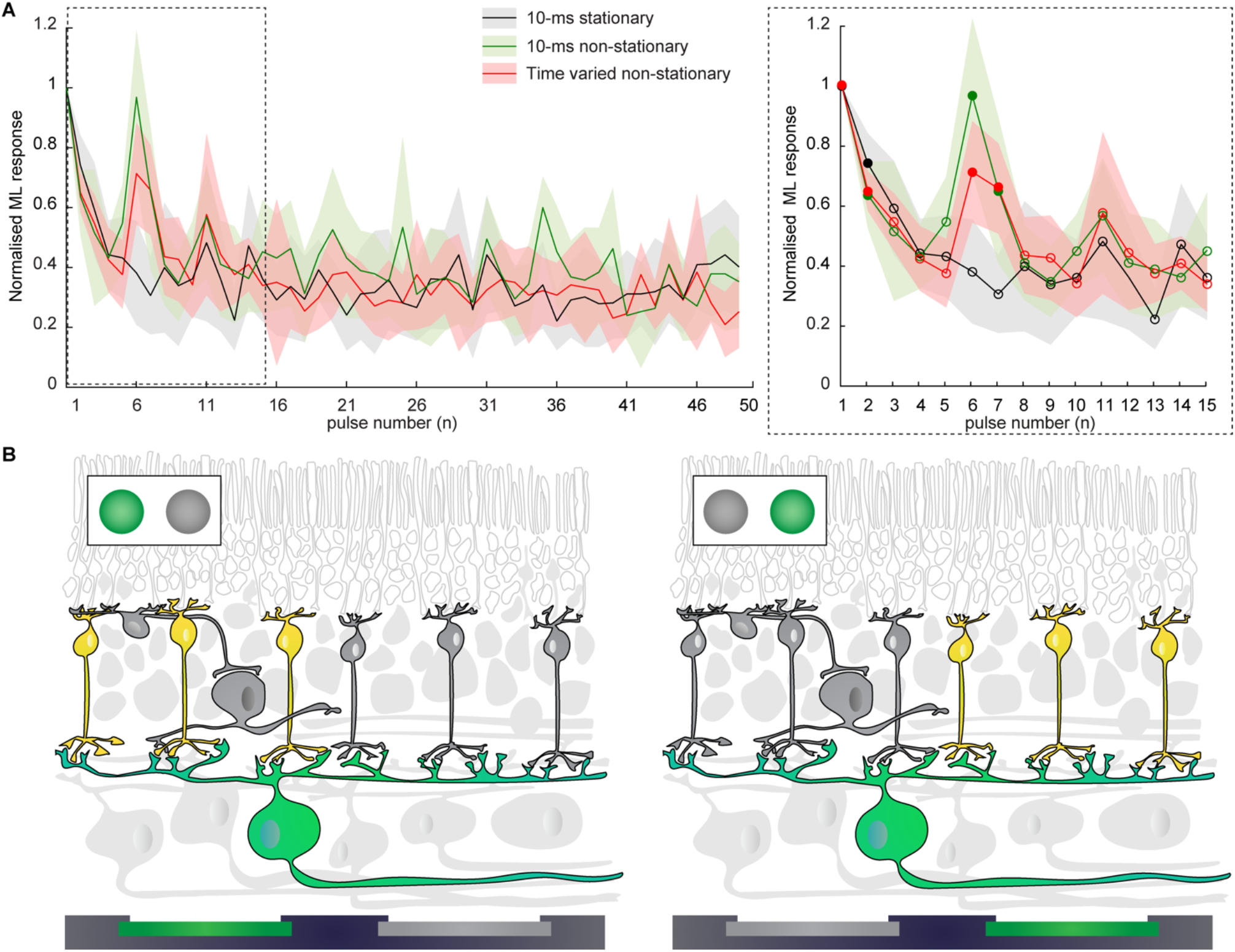
Quantification of the RGC response preservation with non-stationary pulse trains. **A**, Quantification of the normalised medium-latency network-mediated RGC response over 10 s 5-Hz stimulation with 10-ms stationary pulses, 10-ms non-stationary pulses and randomised time-varying non-stationary pulses (mean ± s.e.m., group C, n = 12 RGCs). The firing rates were normalised to the naïve response. The right panel highlights the first 3 s of the recordings. The shaded area shows the s.e.m. Filled circles represent pulses inducing a network-mediated ML activity not statistically significantly different than the naïve repose to the first pulse. Empty circles represent pulses inducing a network-mediated ML activity statistically significantly reduced compared to the naïve repose to the first pulse. **B**, Sub-receptive field stimulation of spatially distinct pools of BCs during non-stationary stimulation. Solid and empty cells represent respectively preserved and degenerated cells in the rd10 mice model. The highlighted cells show the recorded RGC and its presynaptic neurons, among which resting cells are shown in grey, and depolarised cells are coloured.

Non-stationary pulses (both 10-ms and randomised time-varying) lengthened the duration of the RGC response from 0.4 to 1.4 seconds, by reducing the fast RGC desensitisation. According to these findings and previous results [23], we hypothesised that the response recovery at the pixel switch arose from a second pool of BCs that were not desensitised by the first stimulation pattern (Fig. 5B): that is to say that switching the stimulation pixels allows stimulating a RGC alternatively from different subsections of its receptive field.

### Temporal modulation of epiretinal stimulation with interrupted pulse trains

Non-stationary pulses prolonged the persistence of the network-mediated ML response. However, when the illumination spot was switched back to the original pixel, the response was statistically significantly reduced compared to the naïve response. Thus, we introduced another temporal modulation in the stimulation sequence. For each stimulation block (5 pulses at 5 Hz), the last two pulses were removed (interrupted sequence, Fig. 6) since the network-mediated response to these pulses were always statistically significantly reduced compared to the naïve response. Instead, the interrupted sequence leaves a longer gap to each pool of BCs to repolarise. The interrupted sequence design was tested on rd10 explanted retinas (group D in Tab. 1) in combination with 10-ms stationary pulses, randomised time-varying stationary pulses and 10-ms non-stationary pulses (Fig. 6A,B). The interruption of the 10-ms stationary pulse train had no effect since the evoked network-mediated response persisted only for 0.4 (Fig. 6C, blue), similar to 10-ms stationary pulses (Fig. 6C, black). However, the combination of the interrupted sequence with randomised time-varying stationary pulses allowed the persistence of the response up to the third switch (pulse number 16), which corresponds to a response persistence of 3.2 s (Fig. 6C, green). In contrast, the combination of interrupted sequence and 10-ms non-stationary stimulation allowed a response persistence to only 2.2 s (Fig. 6C, yellow).

**Figure 6.**
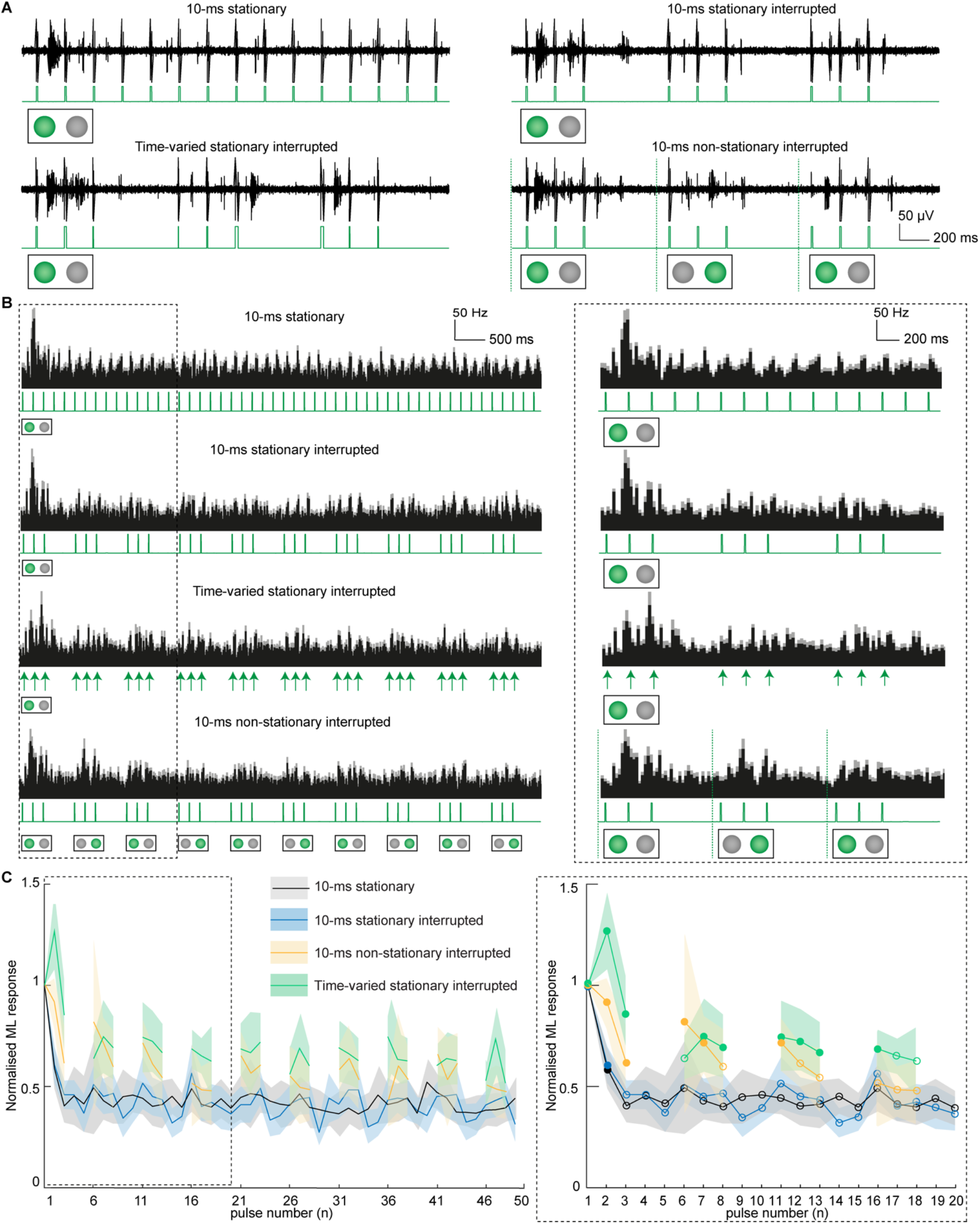
Interrupted pulse trains reduced RGC desensitisation. **A**, Representative recordings from a RGC upon 5-Hz single-pixel photovoltaic stimulation (3 s) with 10-ms stationary pulses (top left), 10-ms stationary interrupted pulses (top right), randomised time-varying stationary interrupted pulses (bottom left) and non-stationary interrupted pulses (bottom right). The light pulses are shown in green. For non-stationary stimulation, the stimulating pixels were switched at 1 Hz. **B**, Mean PSTHs (mean ± s.e.m., group D, n = 23 RGCs) upon 5-Hz single-pixel photovoltaic stimulation (10 s) with 10-ms stationary pulses (first row), 10-ms stationary interrupted pulses (second row), randomised time-varying stationary interrupted pulses (third row) and 10-ms non-stationary interrupted pulses (fourth row). The black area shows the mean value, while the grey area shows the s.e.m. The right panels highlight the first 3 s of the histograms. The light stimuli are indicated in green for 10-ms pulses, while the onset of the light pulses is indicated with green arrows for the randomised time-varying pulses. **C**, Quantification of the normalised network-mediated ML RGC response over 10 s 5-Hz stimulation with 10-ms stationary pulses, 10-ms stationary interrupted pulses, randomised time-varying stationary interrupted pulses and non-stationary interrupted pulses (mean ± s.e.m., group D, n = 23 RGCs). The firing rates were normalised to the response to the naïve pulse. The right panel highlights the first 3 s of the recordings. The shaded area shows the s.e.m. Filled circles represent pulses inducing a network-mediated ML activity not statistically significantly different than the naïve repose to the first pulse. Empty circles represent pulses inducing a network-mediated ML activity statistically significantly reduced compared to the naïve repose to the first pulse.

In summary, randomised time-varying stationary interrupted pulses allows an eightfold lengthening of the RGC response from 0.4 to 3.2 seconds, by strongly reducing the fast RGC desensitisation.

### Spatiotemporal modulation of epiretinal stimulation

Last, we tested on rd10 explanted retinas (group D in Tab. 1) a complex stimulation sequence associating the three modulations described so far (Fig. 7A,B). The combination of non-stationary stimulation together with randomised time-varying pulses and interrupted stimulation sequence theoretically maximise the recovery time between consecutive stimulations of each of the two pools of BCs. Such sequence can reproducibly evoke network-mediated ML responses not statistically different than the first naïve response up to four pixel switches (pulse number 21), corresponding to a response persistence of 4.4 seconds (Fig. 7C).

**Figure 7.**
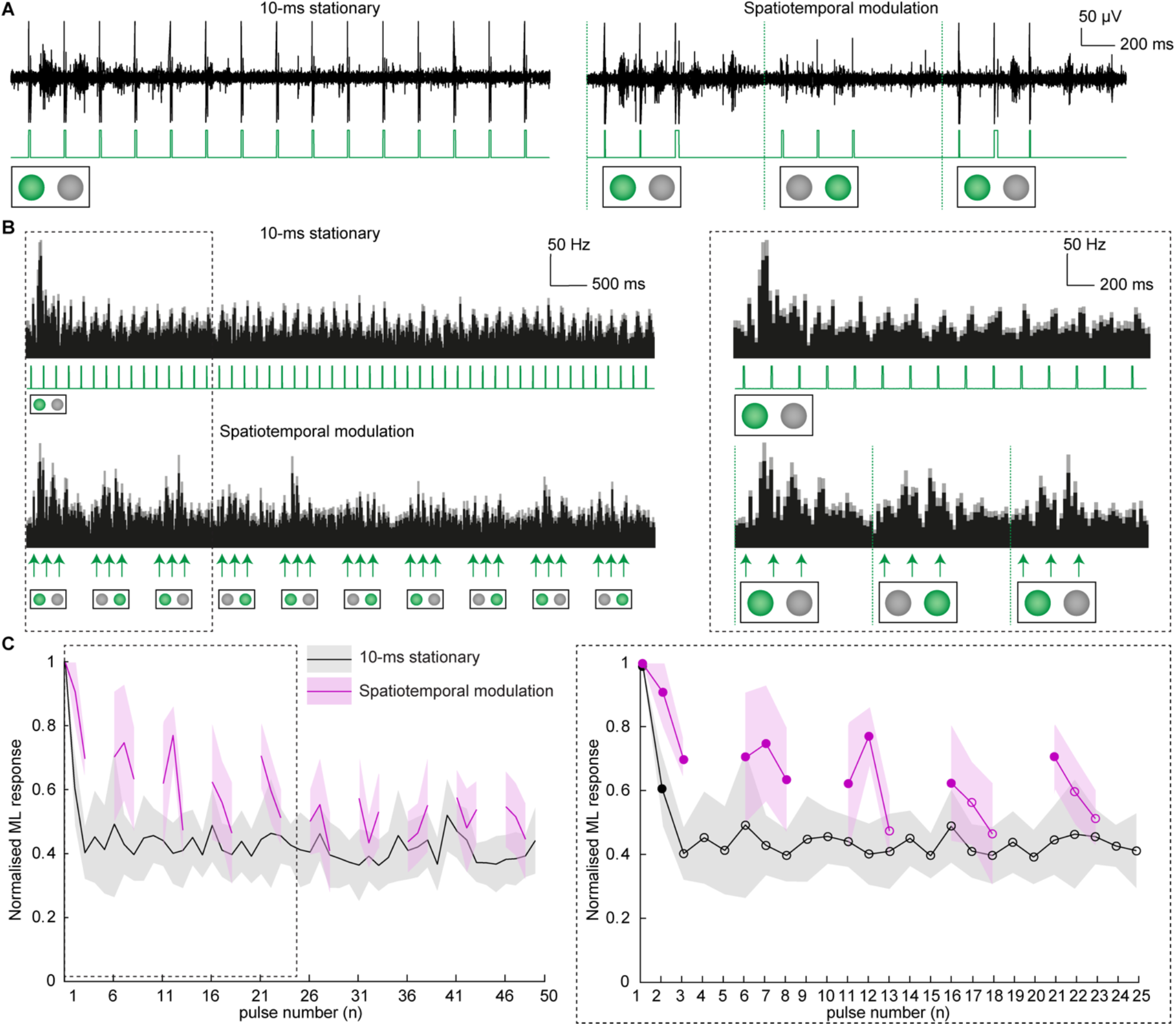
Combined spatiotemporal modulation reduced RGC desensitisation. **A**, Representative recordings from a RGC upon 5-Hz single-pixel photovoltaic stimulation (3 s) with 10-ms stationary pulses (left) and non-stationary randomised time-varying interrupted pulses (right). The light pulses are shown in green. For non-stationary stimulation, the stimulating pixels were switched at 1 Hz. **B**, Mean PSTHs (mean ± s.e.m., group D, n = 23 RGCs) upon 5-Hz single-pixel photovoltaic stimulation (10 s) with 10-ms stationary pulses (left) and non-stationary randomised time-varying interrupted pulses (right). The black area shows the mean values, while the grey area shows the s.e.m. The right panels highlight the first 3 s of the histograms. The light stimuli are indicated in green for 10-ms pulses, while the onset of the light pulses is indicated with green arrows for the randomised time-varying pulses. **C**, Quantification of the normalised network-mediated ML RGC response over 10 s of repetitive stimulation at 5 Hz (mean ± s.e.m., group D, n = 23 RGCs) with 10-ms stationary pulses (back) and non-stationary randomised time-varying interrupted pulses (pink). The firing rates were normalised to the naïve response. The shaded area shows the s.e.m. Filled circles represent pulses inducing a network-mediated ML activity not statistically significantly different than the naïve repose to the first pulse. Empty circles represent pulses inducing a network-mediated ML activity statistically significantly reduced compared to the naïve repose to the first pulse.

When used in conjunction to maximize the recovery time of the excitatory cells, temporal and spatial modulations of the stimulation exhibited a great synergy. A stimulation sequence, whose design included time-varied, non-stationary and interrupted pulse trains, efficiently counteracts the desensitisation process since it offers to each pool of BCs a long recovery period of about 1.8 s to repolarise between consecutive stimulations. Though, the slow RGC desensitisation, likely imputable to inhibitory interneurons, is not affected by those modulations.

## Discussion

### The cognitive burden of transient percepts

The artificial vision provided by retinal implants allows profoundly blind patients to perform visual guided tasks, such as orientation, navigation, object recognition, object manipulation and reading [33,34]. However, the unnaturalistic vision provided by the current retinal prostheses is a significant issue to implanted patients in daily life [18,35,36]. A very small fraction of users decides to keep using their prosthesis at the end of the clinical trial due to the cognitive load of its use [35,37]. This fate matches the abandon rate in cochlear implant or upper-limb prosthesis users that, if not abandoned, they often become an intermittent aid due to both their physiological and cognitive burden [38–41]. Similarly, artificial vision is a highly specific cognitive task, that implies long associative learning but also a permanent multisensory adjustment of the perception [37,35].

In retinal stimulation, electric stimuli are translated into phosphenes organised in a reduced two-dimensional space. The fading of those phosphenes, the absence of depth information and the fragmentation of the visual space make it necessary to adopt multisensorial strategies to localise and recognise the visual cues. Users of retinal prostheses are taught to perform body and head movements to scan the visual scene while performing active viewing through their implant [18]. These movements serve two purposes: to scan a larger portion of the visual field than the one allowed by a small-size implant and to counteract the fading of the phosphenes. Several groups worldwide are developing wide-field implants that could address the first issue from an engineering perspective [42–44]. However, the rapid fading of the percept still forces the users to voluntarily refresh the stimulation pattern on their retina, performing large eye and head movements [26].

Neither in natural vision, stationary images can generate continuous firing of RGCs. However, in sighted humans, the image refresh is accomplished by a variety of micromovements: ocular drifts and microsaccades cause frequent changes in the retinal stimulation allowing to counteract the fading of the neural response [45,46]. Microsaccades are thought to be crucial in this process [21,31,47].

Artificial vision requires the subject to fixate a visual target in order to identify it; therefore, it accounts on the retinal response to one or several stationary electrodes at a time, leaving open the likelihood of response fading, which requires cognitively demanding eye or head movements. It was suggested that when stimulating the retina repetitively and synchronously, a minimal delay of 155 ms (6.5 Hz) was required between stimuli to avoid fading [48]. However, RGC desensitisation was also observed at lower stimulation frequencies [16]: already from 1 Hz in the present study. Another study also argued that repeated stimulation of distant stimulation sites (at least 790 μm) could avoid fading, suggesting that counteracting fading goes through large spatial variations of RGCs targets during fixation [48]. With a high-resolution photovoltaic approach, we showed that two adjacent pixels, separated by 120 μm only, could stimulate the same RGC from different subsections of its receptive field and maintain response up to 4.2 seconds. The ability to maintain a sustained and localised RGC response may drastically reduce the need for active eye or head movements and the constant attention required from varying the RGC target for a static object.

### Spatial and temporal modulations reduce the rapid desensitisation

The stability of the percept evoked by retinal stimulation depends on several factors, among which the electrode location, the stimulation frequency but also the integrity of the retinal network. In rd10 mice retinas, the RGC, the inner nuclear and the plexiform layers are largely preserved at advanced stage of degeneration [49–51], and the time course of RGC desensitisation is not affected by the degeneration process [17]. The integrity of the excitatory and inhibitory pathways is crucial in the fading process. The dissection of the RGC desensitisation into excitatory and inhibitory driven processes allowed for explaining the reports from patients reporting a brisk phosphene fading in hundreds of milliseconds, followed by a low brightness fading out in tens of seconds [18,26]. The nonlinearity we observed in the temporal relationship between desensitisation and stimulation rate corroborates the coexistence of more than one mechanism. Both excitatory and inhibitory interneurons can be activated during electrical stimulation, independently from the electrode location [52,13]. Both the kinetics of BCs voltage-gated Na^+^-channels and the recruitment of the inhibitory network are two valid candidates to explain the decay of the RGC responsivity in epiretinal stimulation [17,53,54]. Growing evidence suggests that AC to BC synapses may be essential players in the slow desensitisation process. Inhibitory postsynaptic potentials were recorded in BCs after depolarising responses to electrical stimulation. As a consequence of the joint presence of inhibitory postsynaptic potentials and Na^+^-channels closure time-course, BCs require from 0.2 to 2 seconds to fully repolarise after a depolarising voltage transient [17]. The spatiotemporal modulations of the stimulation explored in this paper showed the ability to counterbalance the rapid phase of the RGCs desensitisation, playing on the BC excitatory pathway. The common thread behind sequence interruption, randomised time-varied pulses and non-stationary stimulation is to allow longer recovery periods for a given BC or pool of BCs.

The need for BC repolarisation requires to fix the prosthesis stimulation rate well below what is technically achievable by the photovoltaic interface [43]. Argus^®^ II patients preferred stimulation rates between 5 to 10 Hz, mainly because of the fading problem [26,29]. In photovoltaic stimulation, a 5-Hz stimulation rate allows compromising between safe exposure time [23], reduced desensitisation, and continuous image perception. Also, it allows introducing some temporal and spatial modulation leaving a long recovery time between two consecutive stimulation of the same pool of BC.

### Artificial microsaccades

The natural fixational microsaccades may help to avoid phosphenes fading. However, oculomotor reflexes are often impaired in retinitis pigmentosa, and patients show multidirectional involuntary eye movements or nystagmus [55–57]. A non-stationary component within the stimulation strategy itself might then be necessary to reproduce microsaccades and reduce fading. Eye-tracking systems can operate up to 120 Hz, which can effectively gaze-lock the grid of stimulating elements onto the retina [58]. Once gaze-locked, the stimuli can be laterally flickered between two adjacent pixels to mimic microsaccades artificially. Sub-receptive field stimulation allows to do so without affecting the overall pattern of activation of the RGC layer and therefore asking for limited brain adaptation.

The 120-μm separating the high-resolution POLYRETINA prosthesis pixels [23] allows flickering the stimulus within a distance that does not exceed the one of natural microsaccades, estimated from 0.01 ° to 1 ° of visual angle [59,60]. Switching the stimulating pixels at 1 Hz also allows matching the natural fixational saccade occurrence of 1 to 2 per second [59].

In conclusion, providing a naturalistic type of stimulation, where the changes in the RGC activity pattern matches the natural spatiotemporal features, might improve the chances of successful adoption of prosthetic stimulation. At first, increasing the persistence of RGC response up to 4.2 seconds may allow reaching the critical phosphene duration for a sound understanding of the percept. Also, other evidence suggests an active role of non-stationary retina stimulation into shaping and enlarging the cortical receptive fields, which in turn allows targeted saccades and proper allocation of attention during faces recognition or visual exploration tasks [61,62].

## Acknowledgements

This work was supported by École polytechnique fédérale de Lausanne, Medtronic and Fondation Pierre Mercier pour la Science.

## Author contributions

N.A.L.C performed the experiments, analysed data and wrote the manuscript. M.J.I.A.L. fabricated the prosthesis. D.G. designed and led the study, and wrote the manuscript. All the authors read and accepted the manuscript.

## References

[1] Ghezzi D 2020 Translation of a photovoltaic retinal prosthesis Nat Biomed Eng 4 137–8

[2] Hamel C 2006 Retinitis pigmentosa Orphanet journal of rare diseases 1 40

[3] Stone J L, Barlow W E, Humayun M S, Juan E de and Milam A H 1992 Morphometric Analysis of Macular Photoreceptors and Ganglion Cells in Retinas With Retinitis Pigmentosa Archives of Ophthalmology 110 1634–9

[4] Santos A, Humayun, Juan E de, Greenburg R, Marsh M, Klock I and Milam A 1997 Preservation of the inner retina in retinitis pigmentosa. A morphometric analysis. Archives of ophthalmology (Chicago, Ill.: 1960) 115 511–5

[5] Hartong D T, Berson E L and Dryja T P 2006 Retinitis pigmentosa The Lancet 368 1795–809

[6] Jensen R J and Rizzo J F 2008 Activation of retinal ganglion cells in wild-type and rd1 mice through electrical stimulation of the retinal neural network Vision Research 48 1562–8

[7] Boinagrov D, Pangratz-Fuehrer S, Goetz G and Palanker D 2014 Selectivity of direct and network-mediated stimulation of the retinal ganglion cells with epi-, sub- and intraretinal electrodes. Selectivity of direct and network-mediated stimulation of the retinal ganglion cells with epi-, sub- and intraretinal electrodes. 11

[8] Schiefer M A and Grill W M 2006 Sites of Neuronal Excitation by Epiretinal Electrical Stimulation IEEE Trans. Neural Syst. Rehabil. Eng. 14 5–13

[9] Grosberg L E, Ganesan K, Goetz G A, Madugula S S, Bhaskhar N, Fan V, Li P, Hottowy P, Dabrowski W, Sher A, Litke A M, Mitra S and Chichilnisky E 2017 Activation of ganglion cells and axon bundles using epiretinal electrical stimulation J Neurophysiol

[10] Alqahtani A, Abed A A, Guo T, Lovell N H and Dokos S 2017 A continuum model of electrical stimulation of multi-compartmental retinal ganglion cells 2017 39th Annual International Conference of the IEEE Engineering in Medicine and Biology Society (EMBC) 2017 39th Annual International Conference of the IEEE Engineering in Medicine and Biology Society (EMBC) (Seogwipo: IEEE) pp 2716–9

[11] Im M and Fried S I 2015 Spatial properties of network-mediated response of retinal ganglion cells to electric stimulation 2015 7th International IEEE/EMBS Conference on Neural Engineering (NER) (IEEE) pp 256–9

[12] Weitz A C, Nanduri D, Behrend M R, Gonzalez-Calle A, Greenberg R J, Humayun M S, Chow R H and Weiland J D 2015 Improving the spatial resolution of epiretinal implants by increasing stimulus pulse duration Sci Transl Med 7

[13] Chenais N A L, Leccardi M J I A and Ghezzi D 2019 Capacitive-like photovoltaic epiretinal stimulation enhances and narrows the network-mediated activity of retinal ganglion cells by recruiting the lateral inhibitory network J. Neural Eng. 16 066009

[14] Im M and Fried S I 2015 Indirect activation elicits strong correlations between light and electrical responses in ON but not OFF retinal ganglion cells The Journal of physiology 593 3577–96

[15] Jensen R J and Rizzo III J F 2007 Responses of ganglion cells to repetitive electrical stimulation of the retina Journal of neural engineering 4 S1

[16] Freeman D K and Fried S I 2011 Multiple components of ganglion cell desensitization in response to prosthetic stimulation Journal of neural engineering 8 016008

[17] Walston S T, Chow R H and Weiland J D 2018 Direct measurement of bipolar cell responses to electrical stimulation in wholemount mouse retina J. Neural Eng. 15 046003

[18] Stronks H C and Dagnelie G 2014 The functional performance of the Argus II retinal prosthesis Expert review of medical devices 11 23–30

[19] Stingl K, Bartz-Schmidt K U, Besch D, Braun A, Bruckmann A, Gekeler F, Greppmaier U, Hipp S, Hörtdörfer G and Kernstock C 2013 Artificial vision with wirelessly powered subretinal electronic implant alpha-IMS Proceedings of the Royal Society B: Biological Sciences 280 20130077

[20] Stingl K, Bartz-Schmidt K U, Besch D, Chee C K, Cottriall C L, Gekeler F, Groppe M, Jackson T L, MacLaren R E and Koitschev A 2015 Subretinal visual implant alpha IMS– clinical trial interim report Vision research 111 149–60

[21] Martinez-Conde S, Macknik S L, Troncoso X G and Dyar T A 2006 Microsaccades counteract visual fading during fixation Neuron 49 297–305

[22] Davuluri N S and Weiland J D 2014 Time-varying pulse trains limit retinal desensitization caused by continuous electrical stimulation 2014 36th Annual International Conference of the IEEE Engineering in Medicine and Biology Society (IEEE) pp 414–7

[23] Chenais N, Airaghi Leccardi M and Ghezzi D 2020 Single-pixel epiretinal stimulation with a wide-field and high-density retinal prosthesis for artificial vision (Bioengineering)

[24] Quiroga R Q, Nadasdy Z and Ben-Shaul Y 2004 Unsupervised Spike Detection and Sorting with Wavelets and Superparamagnetic Clustering Neural Computation 16 1661–87

[25] Sekirnjak C, Hottowy P, Sher A, Dabrowski W, Litke A M and Chichilnisky E J 2006 Electrical stimulation of mammalian retinal ganglion cells with multielectrode arrays Journal of neurophysiology 95 3311–27

[26] Fornos A P, Sommerhalder J, da Cruz L, Sahel J A, Mohand-Said S, Hafezi F and Pelizzone M 2012 Temporal properties of visual perception on electrical stimulation of the retina Investigative ophthalmology & visual science 53 2720–31

[27] Nikolaev A, Leung K-M, Odermatt B and Lagnado L 2013 Synaptic mechanisms of adaptation and sensitization in the retina Nat Neurosci 16 934–41

[28] Euler T, Haverkamp S, Schubert T and Baden T 2014 Retinal bipolar cells: elementary building blocks of vision Nat Rev Neurosci 15 507–19

[29] Mills J O, Jalil A and Stanga P E 2017 Electronic retinal implants and artificial vision: journey and present Eye 31 1383–98

[30] Ahissar E, Arieli A, Fried M and Bonneh Y 2016 On the possible roles of microsaccades and drifts in visual perception Vision research 118 25–30

[31] Randl K R 2017 Active dynamic vision based on micro-saccades

[32] Buettner R, Baumgartl H and Sauter D 2019 Microsaccades as a Predictor of a User’s Level of Concentration Information Systems and Neuroscience (Springer) pp 173–7

[33] Da Cruz L, Coley B F, Dorn J, Merlini F, Filley E, Christopher P, Chen F K, Wuyyuru V, Sahel J and Stanga P 2013 The Argus II epiretinal prosthesis system allows letter and word reading and long-term function in patients with profound vision loss British Journal of Ophthalmology 97 632–6

[34] Edwards T L, Cottriall C L, Xue K, Simunovic M P, Ramsden J D, Zrenner E and MacLaren R E 2018 Assessment of the electronic retinal implant alpha AMS in restoring vision to blind patients with end-stage retinitis pigmentosa Ophthalmology 125 432–43

[35] Erickson-Davis C and Korzybska H 2020 What do blind people “see” with retinal prostheses? Observations and qualitative reports of epiretinal implant users (Neuroscience)

[36] Weiland J D and Humayun M S 2014 Retinal Prosthesis IEEE Trans. Biomed. Eng. 61 1412–24

[37] Finn A P, Grewal D S and Vajzovic L 2018 Argus II retinal prosthesis system: a review of patient selection criteria, surgical considerations, and post-operative outcomes Clinical ophthalmology (Auckland, NZ) 12 1089

[38] Biddiss E and Chau T 2007 Upper-limb prosthetics: critical factors in device abandonment American journal of physical medicine & rehabilitation 86 977–87

[39] Cajal M 2013 Surdités, implants cochléaires et impasses relationnelles (Eres)

[40] Gourinat V 2015 Le corps prothétique: un corps augmenté? Revue d’éthique et de théologie morale 75–88

[41] Lane F J, Nitsch K P and Scherer M 2016 Ethical considerations in the development of neural prostheses Neurobionics: the Biomedical Engineering of Neural Prostheses 294–318

[42] Ameri H, Ratanapakorn T, Ufer S, Eckhardt H, Humayun M S and Weiland J D 2009 Toward a wide-field retinal prosthesis J. Neural Eng. 6 035002

[43] Ferlauto L, Leccardi M J I A, Chenais N A L, Gilliéron S C A, Vagni P, Bevilacqua M, Wolfensberger T J, Sivula K and Ghezzi D 2018 Design and validation of a foldable and photovoltaic wide-field epiretinal prosthesis Nature Communications 9 992

[44] Lohmann T K, Haiss F, Schaffrath K, Schnitzler A-C, Waschkowski F, Barz C, van der Meer A-M, Werner C, Johnen S, Laube T, Bornfeld N, Mazinani B E, Rößler G, Mokwa W and Walter P 2019 The very large electrode array for retinal stimulation (VLARS)—A concept study J. Neural Eng. 16 066031

[45] Kagan I, Gur M and Snodderly D M 2008 Saccades and drifts differentially modulate neuronal activity in V1: effects of retinal image motion, position, and extraretinal influences Journal of Vision 8 19–19

[46] Greschner M, Bongard M, Rujan P and Ammermüller J 2002 Retinal ganglion cell synchronization by fixational eye movements improves feature estimation Nature neuroscience 5 341–7

[47] Intoy J and Rucci M 2020 Finely tuned eye movements enhance visual acuity Nature communications 11 1–11

[48] Wilke R G, Greppmaier U, Stingl K and Zrenner E 2011 Fading of perception in retinal implants is a function of time and space between sites of stimulation Investigative Ophthalmology & Visual Science 52 458–458

[49] Gargini C, Terzibasi E, Mazzoni F and Strettoi E 2007 Retinal organization in the retinal degeneration 10 (rd10) mutant mouse: A morphological and ERG study J Comp Neurol 500

[50] Mazzoni F, Novelli E and Strettoi E 2008 Retinal ganglion cells survive and maintain normal dendritic morphology in a mouse model of inherited photoreceptor degeneration Journal of Neuroscience 28 14282–92

[51] Jae S A, Ahn K N, Kim J Y, Seo J H, Kim H K and Goo Y S 2013 Electrophysiological and Histologic Evaluation of the Time Course of Retinal Degeneration in the rd10 Mouse Model of Retinitis Pigmentosa Korean J Physiology Pharmacol 17

[52] Jensen R J, Ziv O R and Rizzo J F 2005 Responses of rabbit retinal ganglion cells to electrical stimulation with an epiretinal electrode Journal of neural engineering 2 S16

[53] Hadjinicolaou A E, Savage C O, Apollo N V, Garrett D J, Cloherty S L, Ibbotson M R and O’Brien B J 2014 Optimizing the electrical stimulation of retinal ganglion cells IEEE Transactions on Neural Systems and Rehabilitation Engineering 23 169–78

[54] Hadjinicolaou A E, Meffin H, Maturana M I, Cloherty S L and Ibbotson M R 2015 Prosthetic vision: devices, patient outcomes and retinal research: A review of prosthetic vision research Clin Exp Optom 98 395–410

[55] Cohen E D 2007 Prosthetic interfaces with the visual system: biological issues Journal of neural engineering 4 R14

[56] Schiller P H and Tehovnik E J 2008 Visual prosthesis Perception 37 1529–59

[57] Paraskevoudi N and Pezaris J S 2019 Eye Movement Compensation and Spatial Updating in Visual Prosthetics: Mechanisms, Limitations and Future Directions Front. Syst. Neurosci. 12 73

[58] Wang L, Yang L and Dagnelie G 2008 Virtual Wayfinding Using Simulated Prosthetic Vision in Gaze-locked Viewing: Optometry and Vision Science 85 E1057–63

[59] Engbert R and Mergenthaler K 2006 Microsaccades are triggered by low retinal image slip Proceedings of the National Academy of Sciences 103 7192–7

[60] Rolfs M, Laubrock J and Kliegl R 2009 Microsaccade-induced prolongation of saccade latencies depends on microsaccade amplitude

[61] Tolias A S, Moore T, Smirnakis S M, Tehovnik E J, Siapas A G and Schiller P H 2001 Eye Movements Modulate Visual Receptive Fields of V4 Neurons Neuron 29 757–67

[62] Martinez-Conde S, Otero-Millan J and Macknik S L 2013 The impact of microsaccades on vision: towards a unified theory of saccadic function Nature Reviews Neuroscience 14 83–96

